# Impact of culture medium on linezolid PK/PD activity against *Staphylococcus aureus* in a Hollow-Fiber infection model simulating cerebrospinal fluid exposure

**DOI:** 10.64898/2026.11.16.767216

**Authors:** N. Prébonnaud, A. Chauzy, I. Martin, C. Adier, J. Moreau, C. Dahyot-Fizelier, S. Marchand

## Abstract

This study aimed to assess the impact of the culture medium on the effect of linezolid against *Staphylococcus aureus* using Time-Kill curves (TKC) and Hollow-Fiber infection model (HFIM) by reproducing cerebrospinal fluid (CSF) PK profiles.

TKC were performed over 30 h in Mueller-Hinton Broth (MHB), artificial CSF (aCSF) and human CSF (hCSF) at 0, 0.25, 1, 4 x linezolid MIC. Clinical CSF concentrations after 600 or 900 mg q12h and 900 mg q8h were reproduced in HFIM over 96 h in MHB and aCSF. A PK/PD model developed from MHB TKC and HFIM data was combined with hCSF TKC data to predict bacterial counts in HFIM in hCSF.

In TKC, bacterial growth was slower in hCSF than MHB. Linezolid was bactericidal in MHB, but only bacteriostatic in hCSF. In aCSF, no growth and no decrease in bacterial counts was observed. In HFIM, bacterial growth was predicted to be also slower in hCSF than MHB. Linezolid at 900 mg q8h was needed to obtain a bactericidal effect in MHB, while 600 mg q12h would be sufficient in hCSF. Despite aCSF renewal in HFIM, no growth and no impact of the treatment was observed whatever linezolid dosing regimen.

Bacterial growth and linezolid effect differed between media. MHB overestimated bacterial growth and suggested higher dosing regimen than hCSF, questioning its suitability for central nervous system infections *in vitro* studies. Using aCSF would be a promising alternative, although its composition requires optimisation to better reproduce bacterial growth and linezolid effect observed in hCSF.

## Introduction

*Staphylococcus* spp., particularly *Staphylococcus epidermidis* and *Staphylococcus aureus,* are the most frequent bacteria responsible for healthcare-associated ventriculitis and meningitis (1–3). Vancomycin is recommended as the first-line treatment for these infections, despite low concentrations reached in the cerebrospinal fluid (CSF) due to its limited ability to penetrate brains barriers (blood-brain barrier and blood-CSF barrier) even in case of inflammation (4,5).

Although linezolid was approved by the Food and Drug Administration for the treatment of Gram-positive bacteria such as methicillin-resistant *Staphylococcus aureus* in hospital-acquired pneumonia and complicated skin infections, *Streptococcus pneumoniae* in community-acquired pneumonia and vancomycin-resistant *Enterococcus faecium* in bacteriaemia (6), it is also recommended as an alternative of vancomycin in healthcare-associated ventriculitis and meningitis when its minimal inhibitory concentration (MIC) is higher than 1 mg/L (4).

Indeed, linezolid crosses brains barriers well with a CSF penetration of around 80% (7,8) making it an interesting antibiotic in the treatment of central nervous system (CNS) infections. However, although *Staphylococcus* spp. with MIC ≤ 4 mg/L are consider susceptible to linezolid (9), the recommended dosing regimen of 30-minute infusion of 600 mg twice daily seemed to be insufficient to treat healthcare-associated ventriculitis and meningitis caused by staphylococci with an MIC ≥ 1mg/L, hence the need to optimize the dosing regimen (8,10).

Optimization of dosing regimen can be achieved through *in vitro* pharmacokinetic/pharmacodynamic (PK/PD) studies using the Hollow-Fiber infection model (HFIM) (11). Traditionally, human plasma concentrations are reproduced in HFIM (11–16). However, in the context of CNS infections, where antibiotic diffusion is governed by brain barriers, the PK profiles of antibiotics in the CSF can differ significantly from systemic plasma profiles (8). This highlights the need to reproduce PK profiles at the site of infection in these *in vitro* models (17).

Additionally, most of HFIM are performed in rich medium such as Mueller-Hinton Broth (MHB) (18–21). Indeed, MHB is recommended by EUCAST and CLSI for MIC determination to improve bacterial growth (9,22). However, this medium, composed of casein acid, beef extract and starch, is far removed from physiological media such as CSF, which is mostly composed by ions such as sodium and a low concentration of proteins, and therefore does not reflect the environment of the bacteria at the infection site (23).

As human CSF (hCSF) is difficult to obtain and large volumes are needed to perform HFIM, the use of artificial CSF (aCSF) could be an alternative, although its ability to replace hCSF still need to be evaluated.

In this context, we aimed to evaluate the effect of linezolid against *Staphylococcus aureus* by improving the *in vitro-in vivo* extrapolation of PK/PD models to dosing regimen optimisation, by reproducing CSF PK profiles in HFIM and by assessing the impact of the culture medium on bacterial growth and linezolid effect by comparing MHB, aCSF and hCSF.

## Materials and methods

### 1. Media

Cation-adjusted Mueller-Hinton broth II (MHB) (Merck KGaA, Saint-Quentin-Fallavier, France) was used. Human CSF (hCSF) was collected from intensive care unit (ICU) patients with external ventricular drain (EVD) included in the clinical trial PK-Pop-LCR (PHRCN 16-0501) (8), then sterilized by filtration with 0.2 µm filter after thawing. The compounds used to prepare aCSF were purchased from VWR (Rosny-sous-Bois, France) for CaCl_2_.2H_2_O, KCl, D(+)-Glucose anhydrous, NaH_2_PO_4_, NaHCO_3_; from Carl Roth (Lauterbourg, France) for MgCl_2_ and from Sigma-Aldrich (Merck KGaA, Saint-Quentin-Fallavier, France) for NaCl, Bovine Serum Albumin (BSA) and Minimum Essential Media (MEM) Amino-Acids 100X. Mueller-Hinton Agar (MHA) (Oxoid, ThermoFisher Scientific, Illkirch-Graffenstaden, France) with or without 1% of activated charcoal (Merck KGaA, Saint-Quentin-Fallavier, France) was used for bacterial counts.

#### Preparation of aCSF

Artificial CSF was prepared based on the physiological composition of CSF from human healthy individuals with slight adjustments (23–28). Two solutions were prepared for a better stability: solution A was an acidic solution that stabilizes glucose and solution B was a basic solution that stabilizes bicarbonates (29–31). Solution A was prepared at 20X consisting of 1.62 g of CaCl_2_.2H_2_O, 1.05 g of MgCl_2_, 80.44 g of NaCl, 2.16 g of KCl, 5.40 g of D(+)-Glucose anhydrous and of 98.180 mL of amino-acids. Osmosis water was added QS 500 mL. Solution B was prepared at 50X consisting of 1.72 g of NaH_2_PO_4_.2H_2_O, 6.93 g of NaHCO_3_ and 3 g of BSA to mimic proteins concentration. Osmosis water was added QS 200 mL. Then, solutions A and B were sterilized by filtration with 0.2 µm filter and stored at 4°C for up to 4 weeks (29). Artificial CSF (aCSF) was reconstituted in 465 mL of autoclaved water by adding 25 mL of solution A and 10 mL of solution B and was stored at 4°C for up to 4 weeks (29). The final composition of aCSF was as follows: 1.1 mM of Ca^2+^, Mg^2+^ and H_2_PO_4_^-^, 8.25 mM of HCO_3_^-^, 147 mM of Na^+^, 2.9 mM of K^+^, 145 mM of Cl^-^, 3 mM of glucose, 0.7 mM of amino-acids, 0.3 g/L of BSA, resulting in a pH of approximately 7.3.

### 2. Antimicrobial agents and chemicals

Linezolid standard (purity > 99% in powder form) was purchased from Sigma-Aldrich (Merck KGaA, Saint-Quentin-Fallavier, France). Linezolid D8 (purity of 100% in powder form) was obtained from Alsachim (Illkirch Graffenstaden, France). Linezolid was reconstituted in dimethyl sulfoxide (DMSO, Merck KGaA, Saint-Quentin-Fallavier, France) to obtain stock solutions either at 10,240 mg/L for Time Kill Curves (TKC), or at 10,000 mg/L for HFIM. Stock solutions were prepared at high concentrations to achieve a final DMSO percentage of less than 2% during *in vitro* experiments in order to avoid an antimicrobial effect related to DMSO (32). Stock solutions were stored at -80°C for a maximal duration of 1 month. The day of the experiment, stock solutions were thawed, diluted to the desired concentration either in MHB, hCSF or aCSF for TKC or in 0.9% sodium chloride (Merck KGaA, Saint-Quentin-Fallavier, France) and then transferred in an infusion bag for HFIM.

### 3. Time-Kill curves

TKC were performed on *Staphylococcus aureus* ATCC (*American Type Culture Collection*) 29213 (SA29213) (Linezolid MIC = 4 mg/L) in triplicate in a final volume of 2 mL of MHB, hCSF and aCSF in a 24-well plate (Cellstar®, Greiner, Dutscher, Bernolsheim, France). For all media, linezolid was added to the wells of the plate to obtain concentrations corresponding to 0, 0.25, 1, 4 x MIC. Then, the same steps were followed for all media, with slight adjustments to the MHB protocol for hCSF and aCSF, as described below.

#### MHB

Bacterial strains were cultured overnight at 35 ± 2 °C in 10 mL of MHB under shaking at 130 rpm. The day after, a preculture was performed to achieve exponential growth phase by diluting the overnight culture to 1:50 in a final volume of 10 mL of fresh MHB and incubated at 35 ± 2 °C under shaking at 130 rpm during 1.5 h to 2 h. Optical density was measured at 600 nm with an UltrospecTM 10 Cell Density Meter (Biochrom, USA). Then, the preculture was diluted in fresh MHB to obtain a bacterial suspension at 2 x 10^6^ CFU/mL. A volume of 1 mL was then added to the wells previously filled with 1 mL of MHB supplemented with linezolid, resulting in a final inoculum of 1 x 10^6^ CFU/mL. The plate was incubated at 35 ± 2 °C under shaking at 130 rpm during 30 h.

#### hCSF and aCSF

Bacterial strains were cultured overnight under shaking at 300 rpm in 5 mL of hCSF or aCSF at 35 ± 2 °C with 5% CO_2_. The overnight cultures were diluted in the plate to obtain an inoculum of 1 x 10^6^ CFU/mL in a final volume of 2 mL of aCSF and hCSF. The plate was incubated under shaking at 300 rpm during 30 h at 35 ± 2 °C with 5% CO_2_ to maintain the pH at 7.3 (33).

For all media, a sample of 20 µL was taken from each well at 0, 3, 6, 24 and 30 h, then serially diluted in 0.9% sodium chloride in 96-well plate and plated onto MHA plate to obtain bacterial counts over time. Undiluted samples were plated onto MHA plate supplemented with 1% activated charcoal to avoid linezolid carry-over effect (34–37). The MHA plates were incubated at 35 ± 2 °C during 18 to 24 h, then CFU were counted with a Scan 300. The limit of quantification was fixed to 2.3 log_10_ CFU/mL.

### 4. Hollow-fiber infection model

Typical CSF linezolid concentrations in ICU patients were generated using the clinical population PK model previously described (8). Target CSF concentrations were simulated for three dosing regimens (30-min infusion of 600 mg every 12 h, 30-min infusion of 900 mg every 12 h and 30-min infusion of 900 mg every 8 h) using mrgsolve R package (38) with R software, version 4.2.2 (39). CSF concentrations at these three dosing regimens were reproduced for 96 h in the HFIM in duplicate, on separate occasions, in MHB and aCSF, as previously described (17). SA29213 was inoculated into the extracapillary space (ECS) of a dialysis cartridge (FX paed helixone dialyzer, Fresenius Medical Care, Bad Homburg, Germany) to obtain 1 x 10^6^ CFU/mL. All HFIM were incubated at 37 °C, while those performed in aCSF were supplemented with 5% CO₂ to maintain the pH at 7.3 (33).

Serial samples were taken at various time points from the central reservoir to assess antibiotic concentrations, and from the ECS of the cartridge to assess both antibiotic concentrations, bacterial counts over time. Antibiotic samples were stored at -80°C until assayed by LC-MS/MS.

### 5. LC-MS/MS assay

LC-MS/MS method to assess linezolid concentrations in MHB was previously described (17) and was adapted for aCSF samples as described in (**Text S1**).

### 6. PK/PD model and simulations

As hCSF volume was insufficient to perform HFIM, we sought to predict the bacterial counts (log_10_ CFU/mL) that might have been observed in HFIM conducted in hCSF for 96 h. To this end, bacterial counts (log_10_ CFU/mL) observed during TKC and HFIM in MHB were first analysed simultaneously using a semi-mechanistic PK/PD model. The previously described clinical population PK model (8) was linked to the PD model, with CSF concentrations driving bacterial killing in the HFIM, while constant linezolid concentrations from the TKC wells drove bacterial killing in TKC (**Figure 1**).

**Figure 1.**
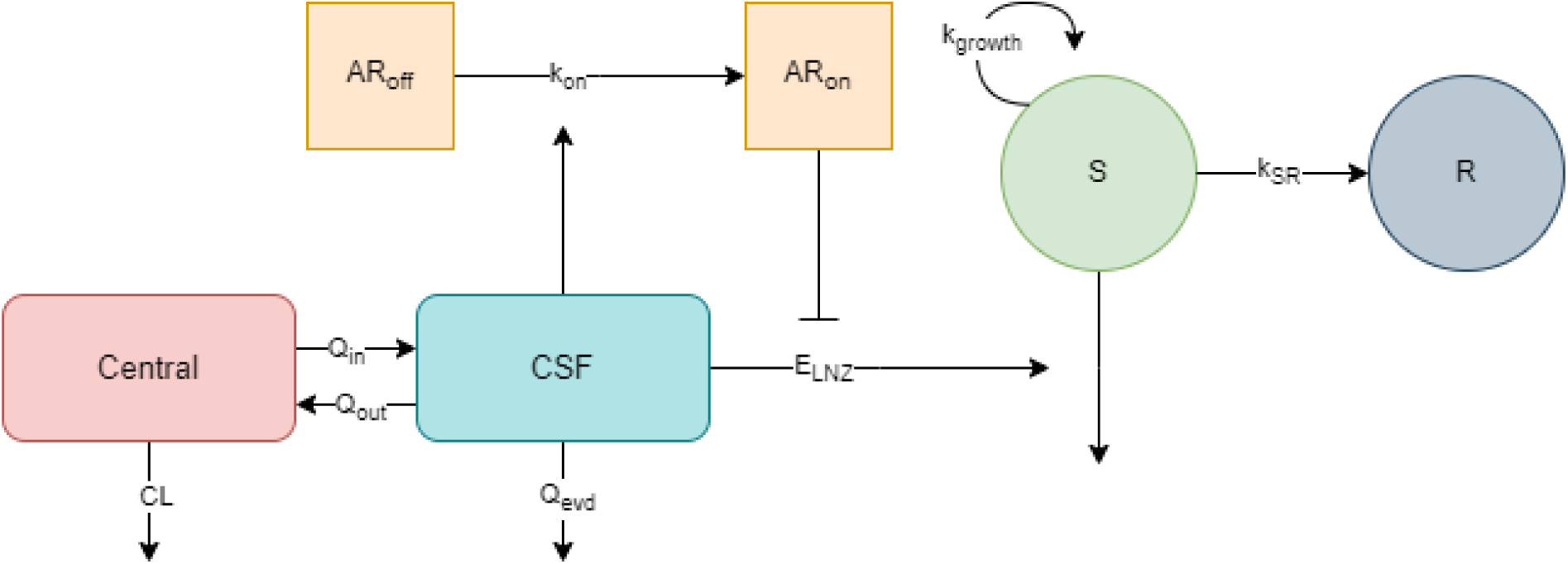
Structure of the PK/PD model. Central: linezolid plasma concentrations (mg/L), CSF: linezolid CSF concentrations (mg/L), CL: clearance (L/h), Q_in_: clearance from central to CSF (L/h), Q_out_: clearance from CSF to central (L/h), Q_evd_: clearance of the external ventricular drain (L/h), AR_off_ and AR_on_: adaptive resistance compartments, k_on_: rate constant for development of adaptive resistance (h^-1^), S: proliferating and susceptible subpopulation (CFU/mL), R: resting and non-susceptible subpopulation (CFU/mL), k_growth_: rate constant for bacterial growth (h^-1^), k_SR_: transfer rate constant from S to R subpopulation (h^-1^), E_LNZ_: linezolid effect (h^-1^).

The PD model consisted of two bacterial subpopulations: one proliferating and susceptible to the antibiotic (S in CFU/mL), and the other resting and non-susceptible (R in CFU/mL). In the starting inoculum, all the bacteria were assumed to be in the growing stage (S), with a first-order rate constant for bacterial growth (k_growth_ in h^-1^). A first-order rate constant for bacterial death (k_death_ in h^-1^) was found to be non-identifiable and was therefore fixed to zero. The transformation of the S subpopulation into the R subpopulation, induced by a high total bacterial level in the system (S + R), was described by a transfer rate constant (k_SR_ in h^-1^) using a linear function (**Eq. 1**), where *B_max_* corresponds to bacterial counts at stationary phase in CFU/mL. However, a reverse transfer from the R subpopulation to the S subpopulation was not considered in the model because the bacteria were continuously exposed to linezolid.

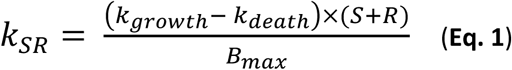

Linezolid was assumed to have an effect (*E_LNZ_*) only on susceptible bacteria and was characterized using an E_max_ function (**Eq. 2**).

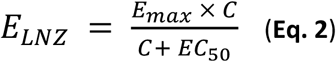

Where E_max_ corresponds to the maximal killing rate due to linezolid in h^-1^, C corresponds directly to the linezolid concentration in mg/L used in the TKC well or to CSF concentration in ICU patients for HFIM and EC_50_ corresponds to the linezolid concentration needed to reach 50% of E_max_ in mg/L.

A function representing adaptation of the bacteria to linezolid was tested, using two states between which bacteria oscillate depending on whether they are adapted (AR_on_) or not (AR_off_). At the beginning of the experiment, all bacteria were assumed to be non-adapted (AR_on_ = 0 and AR_off_ = 1). However, in the presence of linezolid, adaptation developed over time, according to the rate constant k_on_ in h^-1^ leading to a reduction in E_max_ (**Eq. 3**). As bacteria were continuously exposed to linezolid, reverse transfer from the adapted to the non-adapted bacteria was not considered in the model.

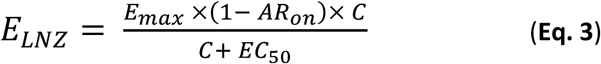

Thus, the development of adaptive and non-adaptive resistance as well as susceptible and non-susceptible bacteria over time were described respectively by **Eq. 4** to **Eq. 7**.

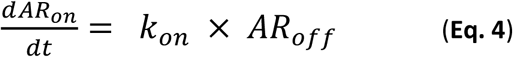

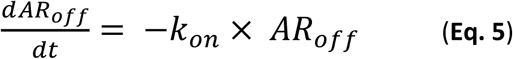

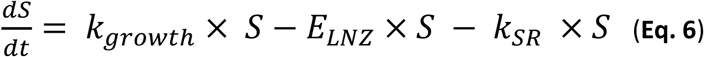

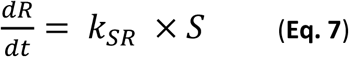

Secondarily, TKC data in hCSF were added and the effect of the medium as a categorical covariate was tested on B_max_, k_growth_, E_max_, EC_50_ and k_on_. PD_parameter_hCSF was then computed using **Eq.8**.

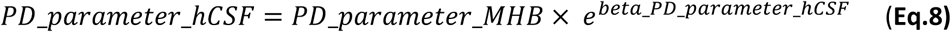

The model was developed using the Stochastic Approximation Expectation-Maximization (SAEM) algorithm in Monolix 2024R1 (40). Bacterial counts data were log_10_-transformed before parameter estimation, and an additive residual error model was applied on the log_10_ scale. Bacterial counts below the LOQ were handled in Monolix based on the M3 method (41).

Model selection was based on goodness of fit plots, objective function value, and relative standard errors (RSE) of the parameter estimates (42). Model fit to data was evaluated by performing visual predictive checks (VPCs) based on 1000 simulations from the final model.

Typical parameter values from the final PK/PD model were used to simulate bacterial counts in hCSF for 96 h after three dosing regimens (30-min infusion of 600 mg every 12 h, 30-min infusion of 900 mg every 12 h and 30-min infusion of 900 mg every 8 h), using the mrgsolve R package (38) with R software version 4.6.0 (39).

## Results

### 1. TKC

Bacterial counts of SA29213 observed during TKC experiments in MHB, hCSF and aCSF are presented in **Figure 2**. In the absence of antibiotic, in MHB, exponential growth was observed until 6 h, followed by a plateau reaching approximately 10^9^-10^10^ CFU/mL, whereas a slower growth was observed in hCSF, with a plateau reaching approximately 10^8^ CFU/mL, and no growth was observed in aCSF. Addition of static linezolid concentrations in MHB showed a concentration-dependent effect, resulting in a decrease of about 4-log_10_ CFU/mL at 30 h at the highest linezolid concentration tested (16 mg/L). In contrast, in hCSF, the maximum effect was less pronounced than in MHB with a decrease of about 1-log_10_ CFU/mL at 30 h for 4 and 16 mg/L of linezolid. In aCSF, no decrease in bacterial counts was observed regardless of the linezolid concentrations added. However, during the first 6 hours, no decrease in bacterial counts was observed whatever linezolid concentrations both in hCSF and aCSF.

**Figure 2.**
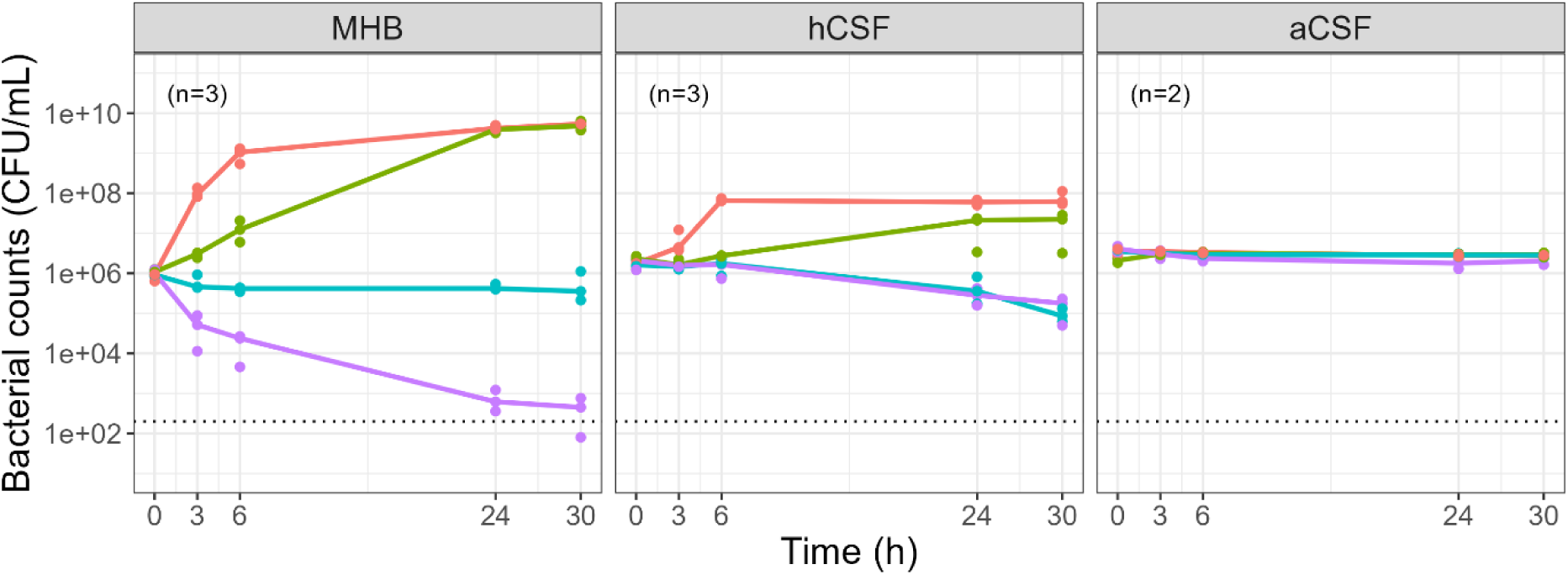
Bacterial counts of SA29213 observed during TKC performed in MHB, hCSF and aCSF. Points correspond to observed bacterial counts for each replicate. Lines correspond to median bacterial counts of the replicates (n) for linezolid concentration of 0 mg/L (red), 1 mg/L (green), 4 mg/L (blue) and 16 mg/L (purple). Dashed line corresponds to the limit of quantification.

### 2. HFIM

#### Pharmacokinetics results

CSF linezolid concentrations were well reproduced in the HFIM performed in MHB and aCSF for all dosing regimens tested with a bias of less than 20% for most concentrations (**Figure 3**).

**Figure 3.**
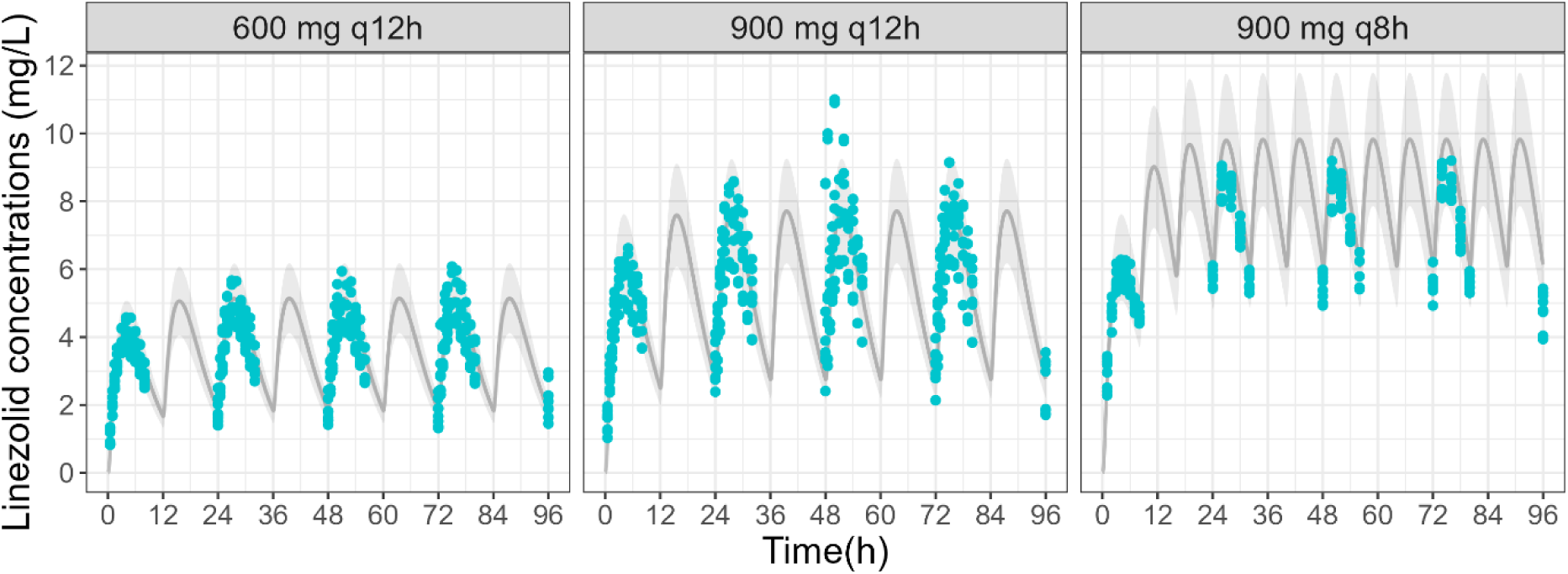
Targeted linezolid CSF concentrations and observed linezolid concentrations in the central reservoir and the ECS of the cartridge of HFIM performed in MHB and aCSF after infusion of 600 mg q12 h, 900 mg q12 h and 900 mg q8 h *(n = 4)*. The grey line corresponds to targeted linezolid concentrations from linezolid population PK model (8). The grey area corresponds to a 20% bias from targeted linezolid concentrations. Points correspond to linezolid concentrations observed in the central reservoir and the ECS of the cartridge of the HFIM in MHB (yellow) and in aCSF (blue).

#### Pharmacodynamics results

Bacterial counts of SA29213 observed in MHB compared to those observed in aCSF in the HFIM are presented in **Figure 4**.

**Figure 4.**
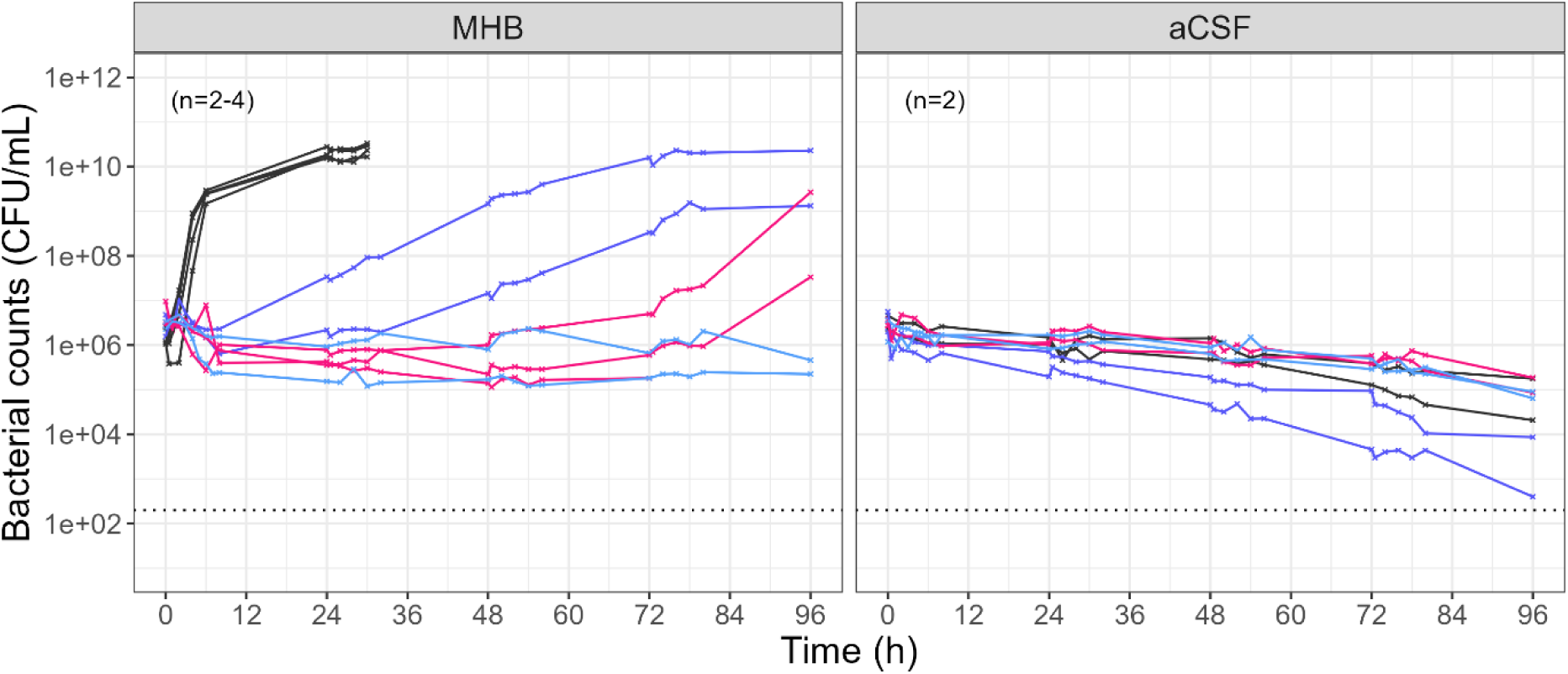
Bacterial counts of SA29213 observed in HFIM performed in MHB and in aCSF for three dosing regimens of linezolid. After a 30-min infusion of 600 mg every 12 h (purple), a 30-min infusion of 900 mg every 12 h (pink), a 30-min infusion of 900 mg every 8 h (blue) of linezolid and in the absence of linezolid (black). In MHB, in the absence of linezolid, bacterial counts were stopped at 30 h due to the presence of the bacteria studied in the central reservoir after 30 h.

In MHB, in absence of linezolid, bacteria exhibited exponential growth reaching ≈ 10^10^ CFU/mL from 24 h, as observed in TKC. The control experiment performed in MHB was repeated 4 times and stopped at 30 h due to the presence of the studied bacteria in the central reservoir (and consequently in the waste) of the HFIM. After confirming that it was not a contamination, it appeared that the bacteria could cross the pores of the capillaries in the cartridge only with a high bacterial inoculum and in the absence of linezolid, as this was not observed under other tested conditions. In MHB, for the recommended dosing regimen of a 30-min infusion of 600 mg every 12 h, no bacterial killing was observed but bacterial growth was slower than in control experiments. Consequently, the bacterial count remained stable during the first 8 h, followed by a growth phase until a plateau of approximately 10^10^ CFU/mL was reached between 72 and 96 h. After a 30-min infusion of 900 mg every 12 h of linezolid in MHB, bacterial counts remained stable up to 72 h, but then increased to approximately 10^8^ CFU/mL at 96 h. After a 30-min infusion of 900 mg every 8 h, bacterial growth was totally inhibited, with bacterial counts remaining stable at approximately the inoculum level (10^6^ CFU/mL) throughout the entire experiment.

In aCSF, despite medium renewal in the HFIM, no bacterial growth was observed at 30h, as previously described in the TKC. Furthermore, a slight bacterial decrease was observed at 96h. Addition of linezolid, whichever dosing regimen, resulted in the same profile as the control.

### 3. PK/PD model

The final PK/PD model well described TKC and HFIM data in MHB as well as TKC in hCSF as indicated by the VPCs in **Figure 5**.

**Figure 5.**
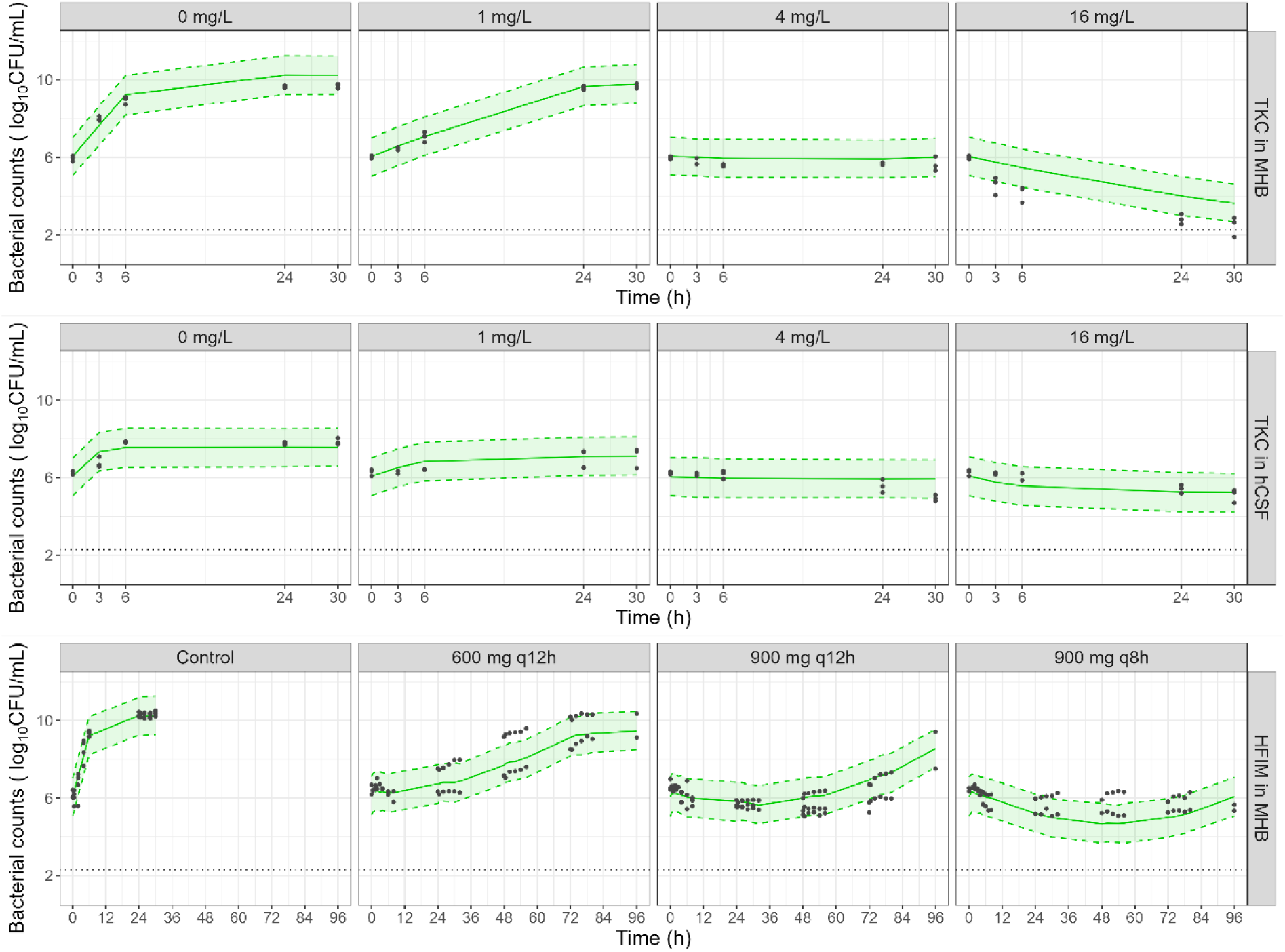
Visual predictive checks (VPCs) for the final PK/PD model. The grey points represent observed bacterial counts (log_10_ CFU/mL). The straight line corresponding to the median of simulations (50^th^ percentile), the dashed lines corresponding to the 5^th^ and 95^th^ percentiles, and the shaded area corresponding to the 90% prediction interval. Dotted black line corresponds to the limit of quantification.

Estimated parameters and RSEs are presented in **Table 1**. The addition of hCSF TKC data required the implementation of a medium effect on B_max_ only. The estimated B_max_ in hCSF was 7.28 log_10_ CFU/mL, which corresponds to a reduction of approximately 3-log compared to the value estimated in MHB.

**Table 1.**
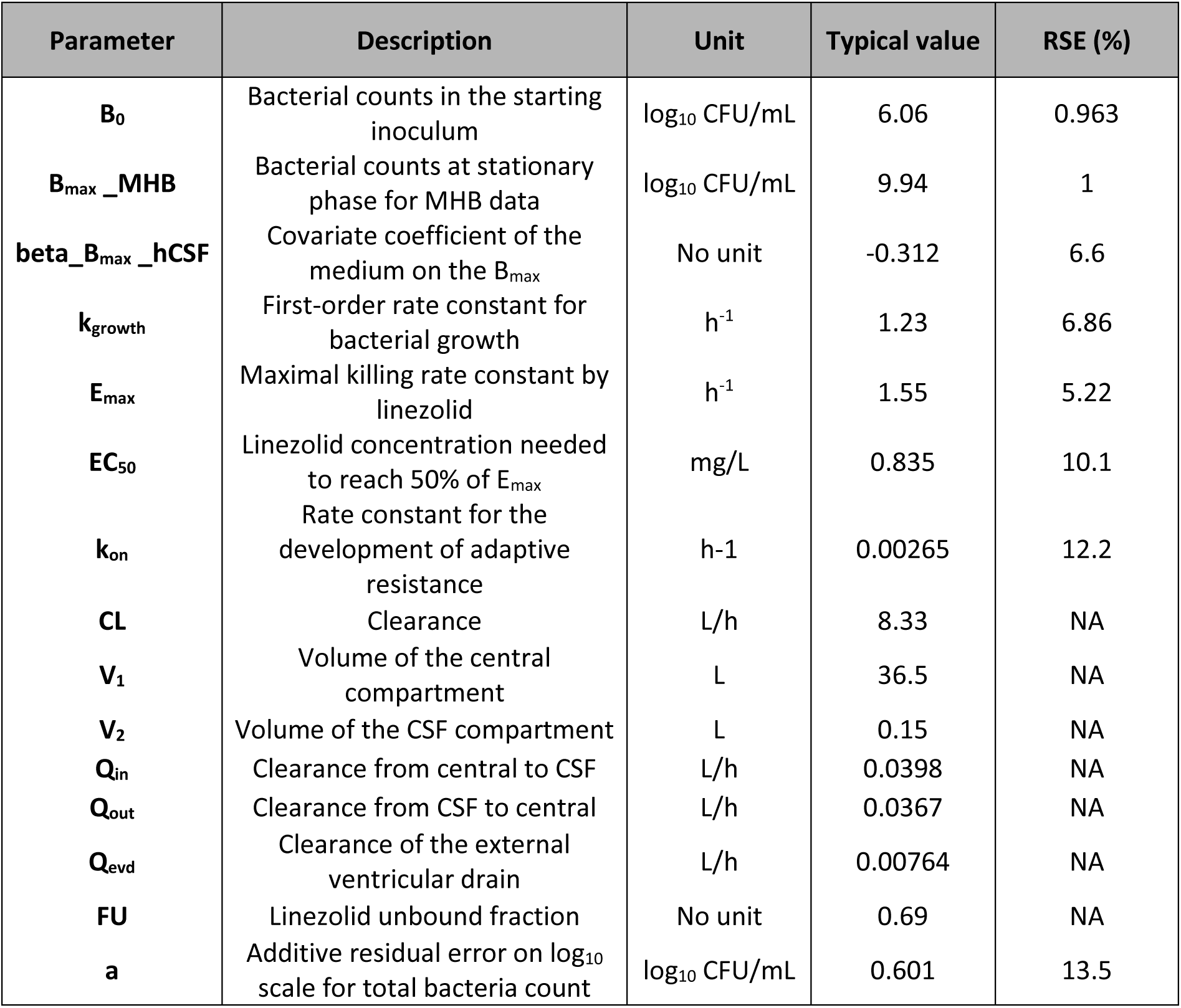
Estimated parameters and relative standard error (RSE). NA: not applicable, as parameters were fixed to the clinical PK model values.

### 4. PK/PD simulations

Predicted bacterial counts of SA29213 in hCSF during HFIM are presented in **Figure 6**.

**Figure 6.**
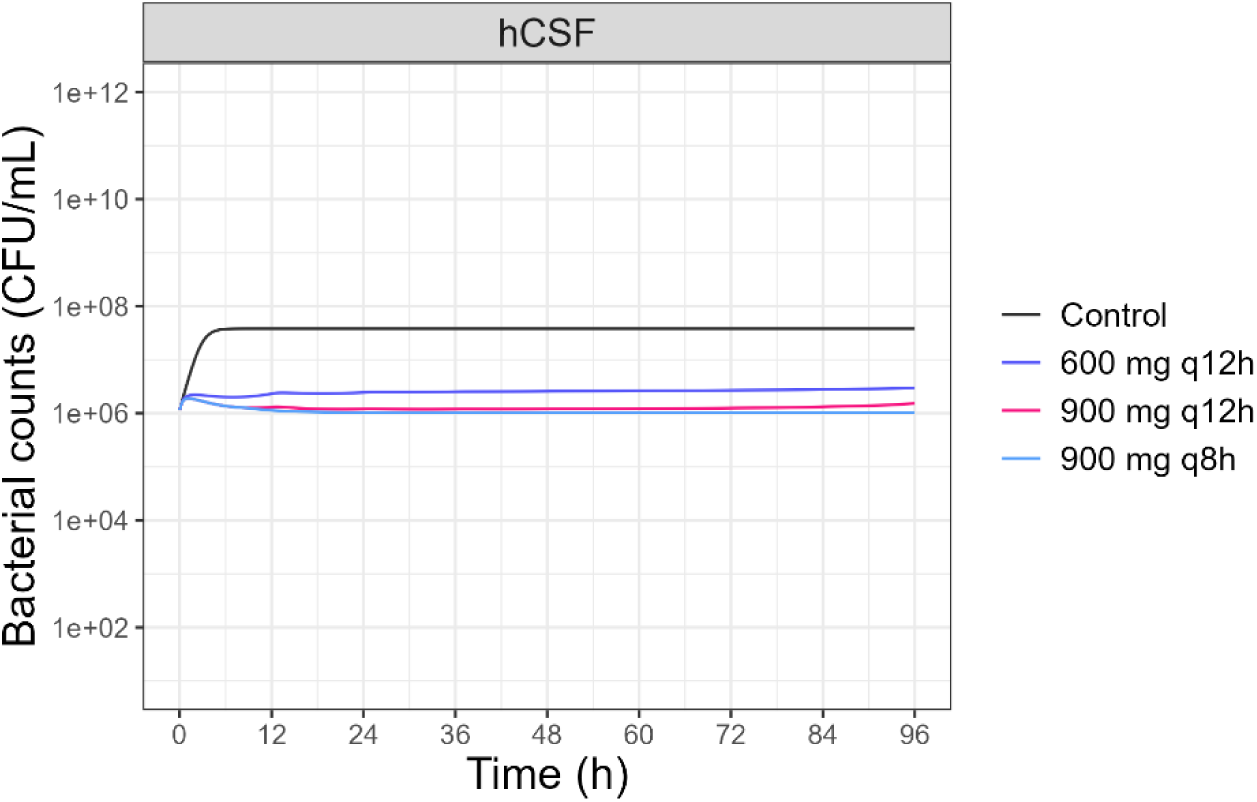
Predicted bacterial counts of SA29213 in HFIM in hCSF for three dosing regimens of linezolid. In the absence of linezolid (black), after a 30-min infusion of 600 mg every 12 h (purple), a 30-min infusion of 900 mg every 12 h (pink), a 30-min infusion of 900 mg every 8 h (blue) of linezolid.

In hCSF, in absence of linezolid, bacteria would exhibit exponential growth reaching ≈ 10^8^ CFU/mL from 6 h. The three dosing regimens would provide a bacteriostatic effect, resulting in inhibition of the bacterial growth with bacterial counts remaining globally stable at the inoculum level (10^6^ CFU/mL) throughout the entire experiment, although the bacteriostatic effect would be slightly lower for 600 mg every 12 h than 900 mg every 12 h or 8 h.

## Discussion

To our knowledge, this is the first study in which linezolid effect against *Staphylococcus aureus* was evaluate using concentrations observed at the infection site, with CSF concentrations reproduced in the HFIM, and the impact of the culture medium investigated by using both MHB and aCSF.

The use of aCSF was intended to provide a standardized medium as an alternative to hCSF. To limit variability, we chose to reproduce the physiological composition of CSF, which remains relatively constant in healthy adults. In contrast, CSF composition may be altered in patients with healthcare-associated ventriculitis and meningitis, depending on the causative microorganisms and the severity of inflammatory response. For instance, in a study of 71 cases of healthcare-associated meningitis, infections caused by coagulase-negative staphylococci were associated with lower proteinorachia and higher glycorrhachia than *Staphylococcus aureus, Enterococcus* spp.*, Streptococcus* spp., *Enterobacteriaceae*, and *Pseudomonas aeruginosa* (43). Additionally, the neurosurgical procedure preceding these infections may themselves induce inflammation, leading to increased proteinorachia (4,44). Consequently, tailoring the composition of aCSF to the microorganism under study and the associated inflammatory response would limit its applicability as a standardized alternative to hCSF.

A previous study aimed to use aCSF as an alternative to hCSF to study the effects of linezolid on *Staphylococcus aureus* and *Staphylococcus epidermidis.* However, the pH of the prepared aCSF was unstable, varying from 7.6 to 5.3, which was unphysiological and, according to the authors, lead to nonsignificant results (45). Unlike the aCSF used in that study (45), we added bicarbonate and sodium dihydrogen phosphate dihydrate to our aCSF and use it in an atmosphere containing 5% CO_2_ to improve pH stability. However, to maintain a pH of 7.3 throughout experiments, the bicarbonate concentration in our aCSF was intentionally reduced compared to that of healthy hCSF as the physiological concentration led to a pH of approximately 8, which was deemed unacceptable. In fact, previous studies have shown that pH, whether acidic or basic, has a strain-specific effect on bacterial growth (46). For example, the growth of *Staphylococcus aureus* was reduced when the pH decreased from 7 to 5, whereas the growth of *Staphylococcus epidermidis* was not affected by this change (47). Additionally, the chloride concentration in our aCSF was higher than the physiological concentration targeted (145 mM vs 113 mM). This difference can be attributed to the presence of chloride ions from other components in the aCSF, making it difficult to reduce the overall chloride concentration without affecting the concentrations of other compounds.

In the present study, the main observation is that growth profiles differed between media, with bacterial growth being slower in both hCSF and aCSF than in MHB (**Figure 2**). This observation is consistent with the findings of Agbayani *et al*. who compared the bacterial growth of five bacterial species, including *Staphylococcus aureus* in trypticase soy broth (TSB) and hCSF. Although the growth curves were performed under aerobic conditions, raising concerns about potential pH changes and their impact on bacterial growth, slower growth was observed for all the fives bacterial species in hCSF than in TSB (48). This slowdown in growth was also observed during growth control experiments in diluted MHB, from 100% to 1%, on a reference strain of *Pseudomonas aeruginosa* (49), as well as in peritoneal dialysis fluids on reference strains of *Escherichia coli, Staphylococcus aureus*, *Staphylococcus epidermidis*, *Pseudomonas aeruginosa*, *Enterococcus faecalis* and *Candida albicans,* and was attributed to the less-nutrients composition of these media compared to MHB (50,51).

Furthermore, in our study, aCSF failed to reproduce the bacterial growth observed in hCSF (**Figure 2**), and renewing aCSF in the HFIM did not promote bacterial growth (**Figure 4**), suggesting that the aCSF formulation itself may be insufficient to support bacterial growth. In addition, the hCSF used in this study was collected from ICU patients with EVD, and its composition may differ from that of physiological CSF. These differences suggest that one or more components required for bacterial growth may be missing or insufficiently represented in the aCSF formulation. For instance, glucose concentration may be insufficient, as its serves as a nutrient source for bacteria. However, a previous study showed that increasing glucose concentrations in peritoneal dialysis fluids did not enhance *S.aureus* growth, suggesting that the non-growing state observed in our study may not due to a lack of glucose (50,51). Moreover, the authors reported slightly better bacterial growth with the use of peritoneal fluids containing a mix of 15 amino acids than in the absence (50,51). Unlike other aCSF formulations, our aCSF contains amino acids (26,29–31,45,52–54), but only a blend of 7 have been added. Among them, 4 amino acids (glutamic acid, aspartic acid, proline and alanine) have been reported as key amino acids during the mid-exponential growth of both clinical and reference strains of *Staphylococcus aureus* (55). However, in this study, they detected 22 amino acids in the cytoplasm of these bacteria during mid-exponential growth, with variations depending on the reference and clinical strains (55), suggesting that additional supplementation with other amino acids may be an option to enhance bacterial growth in aCSF.

In addition to the differences in growth profiles, the effect of linezolid also varied according to the medium. In the present study, linezolid exhibited a bactericidal effect in MHB, while it was only bacteriostatic in hCSF (**Figure 2**). Our results partially align with those reported by Schwameis *et al*. (45). Indeed, although linezolid caused less bacterial decrease in MHB (*i.e*. bacteriostatic effect at 4 times the MIC), their profiles in hCSF remained consistent with ours, showing a bacteriostatic effect whatever the linezolid concentration (45). This difference in antibacterial activity was also observed in Gram-negative bacteria. Based on the ratio of minimum bactericidal concentration (MBC) to MIC, cefotaxime and piperacillin exhibiting a bacteriostatic effect (MBC/MIC ratio > 4) on *Morganella morganii* in hCSF, whereas both antibiotics exhibited a bactericidal effect (MBC/MIC ratio ≤ 4) in MHB, even though the MIC was lower in CSF than in MHB (56). However, in aCSF, as the bacteria appear to be in a non-growing state, no impact of linezolid was observed (**Figure 2**). This poor artificial medium seems to activate a stress response that inhibits division and could be responsible for a tolerance phenomenon in the *Staphylococcus aureus* strain (57). In this state, bacterial protein synthesis may be inactive, rendering linezolid ineffective, as its mechanism of action (blocking the initiation of translation) targets a process that may already dormant. A similar mechanism may also occur in hCSF. Indeed, when hCSF TKC data were included into the PK/PD model, a covariate effect on B_max_ alone was required to account for the effect of the medium. This resulted in a lower B_max_ in hCSF and, according to **Eq. 1**, a higher k_SR_, leading to an increase in the R population, which represents a non-growing bacterial state that is insensitive to antibiotics. To confirm this hypothesis, additional experiments, including quantitative RT-PCR or flow cytometry (57), could be performed.

Dynamic HFIM is a powerful *in vitro* tool for studying antibiotic efficacy and optimizing new dosing strategies. While such studies are traditionally conducted in rich culture media, our results suggest that dose adjustments may vary depending on the type of medium used. In fact, dynamic HFIM experiments conducted in MHB (**Figure 4**) suggest that a higher dose than currently recommended (600 mg q12h) would be required, as bacterial growth was prevented only for 900 mg q8 h. Although our HFIM results in aCSF do not allow us to draw a formal conclusion at this stage and the composition of aCSF should be improved, the TKC and HFIM data obtained in MHB, combined with the TKC data obtained in hCSF, enabled us to perform PK/PD simulations to predict bacterial counts that might have been observed during HFIM in hCSF. This approach was made possible as the same PK/PD model was able to describe both TKC and HFIM data observed in MHB, without requiring a covariate to account for differences in experimental setup. This finding is consistent with Nielsen *et al*., who showed that a PK/PD model with parameters estimates based on TKC data could reasonably predict bacterial counts in HFIM (58). Building on this approach, we used TKC data in hCSF to predict bacterial counts that would be expected in HFIM in hCSF, thereby providing insights that would otherwise have been difficult to generate experimentally given the difficulty to obtain hCSF. In contrast to MHB, PK/PD simulations predicted a bacteriostatic effect with all three dosing regimens, suggesting that the currently recommended dosing regimen of 600 mg q 12h may be sufficient to achieve a bacteriostatic effect against *Staphylococcus aureus* in CSF (**Figure 6**). These discrepancy between media highlights the importance of investigating the *in vitro* PK/PD of antibiotics in medium that reflect the target infection site, particularly in the case of CNS infections.

One limitation of this study is that the efficacy of linezolid in different media was only evaluated on a reference strain of *Staphylococcus aureus*. The efficacy of linezolid should be investigated in other *Staphylococcus* spp. strains, both reference and clinical strains, for greater clinical relevance. Another limitation is that this study was only conducted on Gram-positive bacteria, limiting the generalizability of these findings regarding the impact of the culture medium on Gram-negative bacteria. This is particularly noteworthy, as previous studies on *Acinetobacter baumannii* and *Escherichia coli* in diluted MHB and hCSF, respectively, showed that their growth was less affected by nutrient-depleted media (48,59). Moreover, a higher antimicrobial effect was observed in diluted MHB for polymyxin B and meropenem on *Acinetobacter baumanii* and *Pseudomonas aeruginosa*, respectively, and for fosfomycin on *Pseudomonas aeruginosa* in peritoneal dialysis fluid (49,50,59). This work should be complemented by further experiments on Gram-negative bacteria.

To conclude, bacterial growth and linezolid effect in MHB were overestimated compared to hCSF, raising questions about the relevance of using MHB in *in vitro* experiments in the context of CNS infections. Using MHB could lead to increasing the dose to be more effective but that might not be necessary. Therefore, the use of aCSF seemed to be a good alternative, although further improvements in composition are needed to better mimic the growth and the linezolid effect observed in hCSF. Additionally, these experiments should be replicated on other strains, as linezolid effect was studied on only one reference strain of *Staphylococcus aureus*.

## Acknowledgements

The authors thank Agnès Audurier and Laure Prouvensier for their technical assistance and Frederic Tewes, Hélène Mirfendereski and Jonathan Clarhault for their advice concerning the preparation of the artificial cerebrospinal fluid. We thank the patients and their relatives and caregivers, local site clinical and research staff, regional coordinating center staff of the university hospital of Poitiers, and all the members of the PK-Pop-LCR Study for their active participation.

The Shimadzu high-performance liquid chromatography system module (Nexera XR; Shimadzu, Marne la Vallée, France) coupled with a TQ3500 mass spectrometer (Sciex, Les Ulis, France) was obtained through the support of the Nouvelle Aquitaine CPER programs.

This work was supported by the French National Research Agency (ANR), under CAPOEIRA grant agreement number ANR-23-CE17-0008-01.

## Supplementary data

### Text S1: LC-MS/MS assay of aCSF samples

For aCSF samples, the method used to assess CSF concentrations was adapted from the method used previously (8). After ensuring that no matrix effect existed between NaCl and CSF matrices, calibration curves were established in NaCl over 0.1 to 50 µg/mL and quality controls (QC) were prepared in aCSF. Samples were precipitated with 200 µL of H_2_O/MeOH (50/50, v/v) containing the internal standard (Linezolid D8) at 500 ng/mL and vortexed for 5 s. Then, samples were centrifuged at 14 000 rpm for 20 min at 4 °C and 100 µL of supernatant was transferred into a glass vial containing 100 µL of acetonitrile/formate d’ammonium 10 mM (50/50, v/v). A volume of 1 µL was injected.

The system included a Shimadzu high-performance liquid chromatography system module (Nexera XR; Shimadzu, Marne la Vallée, France) coupled with a TQ3500 mass spectrometer (Sciex, Les Ulis, France). The compound was analyzed on an XBridge Peptide BEH300 C18 column (5μm, 2.1 x 150 mm, Waters). The mobile phase consisted of a mixture (50/50, v/v) of 10 mM ammonium formate and acetonitrile delivered isocratically at 0.20 mL/min. Electrospray ionization in positive mode was used for detection. Ions were analyzed in the multiple reaction monitoring, and the following transitions were inspected: m/z 338→296 and 346→304 for linezolid and internal standard, respectively.

The intraday variability was characterized at three concentrations levels (0.3, 5 and 35 µg/mL) with a precision < 4% and bias < 9% (n = 3). The interday variability was characterized at three concentrations levels (0.3, 5 and 35 µg/mL) with a precision < 4% and bias < 3% (n = 9).

